# Cytokit: A single-cell analysis toolkit for high dimensional fluorescent microscopy imaging

**DOI:** 10.1101/460980

**Authors:** Eric Czech, Bulent Arman Aksoy, Pinar Aksoy, Jeff Hammerbacher

## Abstract

**Background:** Multiplexed *in-situ* fluorescent imaging offers several advantages over single-cell assays that do not preserve the spatial characteristics of biological samples. This spatial information, in addition to morphological properties and extensive intracellular or surface marker profiling, comprise promising avenues for rapid advancements in the understanding of disease progression and diagnosis. As protocols for conducting such imaging experiments continue to improve, it is the intent of this study to provide and validate software for processing the large quantity of associated data in kind.

**Results:** Cytokit offers *(i)* an end-to-end, GPU-accelerated image processing pipeline; (ii) efficient input/output (I/O) strategies for operations specific to high dimensional microscopy; and (iii) an interactive user interface for cross filtering of spatial, graphical, expression, and morphological cell properties within the 100+ GB image datasets common to multiplexed immunofluorescence. Image processing operations supported in Cytokit are generally sourced from existing deep learning models or are at least in part adapted from open source packages to run in a single or multi-GPU environment. The efficacy of these operations is demonstrated through several imaging experiments that pair Cytokit results with those from an independent but comparable assay. A further validation also demonstrates that previously published results can be reproduced from a publicly available multiplexed image dataset.

**Conclusion:** Cytokit is a collection of open source tools for quantifying and analyzing properties of individual cells in large fluorescent microscopy datasets that are often, but not necessarily, generated from multiplexed antibody labeling protocols over many fields of view or time periods. This project is best suited to bioinformaticians or other technical users that wish to analyze such data in a batch-oriented, high-throughput setting. All source code, documentation, and data generated for this article are available under the Apache License 2.0 at https://github.com/hammerlab/cytokit.

## Background

Molecular profiling of cell culture and tissue samples traditionally relies on techniques that do not support a diverse panel of protein targets without disturbing important *in situ* characteristics of cells. Immunofluorescence imaging preserves these characteristics but is limited to a small number of expression measurements due to the need to avoid overlapping fluorophore emission spectra. This limitation can be overcome to an extent through repeated imaging of the same specimen over several cycles, where each cycle typically involves capturing images of 3 or 4 markers at a time, however the incubation period necessary between cycles is often hours or days and methods for removing markers from previous cycles can be detrimental to assay quality. By contrast, techniques like Mass Cytometry [1] and Multispectral Flow Cytometry [2] enable the measurement of more target compounds but provide little to no morphological or spatial information. Other methods such as Multiplexed Immunohistochemistry [3] and Multiplexed Ion Beam Imaging [4] overcome these limitations but require special appliances that are not compatible with standard or commercial microscopy platforms. For these reasons, analysis of data from multiplexed fluorescent labeling methods are appealing as they are economical, can be conducted with any fluorescent imaging platform, and rely on well documented immunostaining protocols.

Methods developed for multiplexed fluorescent labeling include Co-Detection by Indexing (CODEX) [5], DNA Exchange Imaging (DEI) [6], and t-CyCIF [7], all of which outline a cyclical protocol in which 2 or 3 antigen markers and a DNA stain are introduced in each cycle prior to imaging. Unlike in the t-CyCIF protocol, where a relatively lengthy antibody incubation step is required as a part of each cycle, both CODEX and DEI begin with a single incubation step in which a large (possibly >100) number of oligonucleotide-conjugated antibodies bind to cognate antigens. Fluorophores bound to complementary oligonucleotide sequences are then introduced to enable the collection of a small number (typically 4) of fluorescent images before being washed away and allowing the process to repeat. These procedures are capable of measuring the expression of tens or hundreds of different proteins but introduce several key computational challenges that make employing them difficult. The primary challenge is that incorporating the larger number of expression targets within 3D image volumes that encompass hundreds of thousands of cells, a sample size often necessary for studying less common cell types [8], leads to 100+ GB raw image sets including well over 10,000 images. As this data is often the product of commodity imaging platforms, substantial processing is necessary for analysis and the time associated with this processing becomes prohibitive for high dimensional acquisitions without GPU acceleration. With GPU acceleration however, we show how a CODEX experiment, shared by Goltsev et al., including 51k 16-bit images (129 GB), 54 expression markers, and ~70k individual cells can be aligned, deconvolved, and segmented for analysis in less than 90 minutes per workstation. Another challenge prevalent among such large imaging assays is managing the heterogeneity that arises across replicates and treatment groups for experiments that span multiple days or weeks and involve multiple laboratory practitioners. Images collected as a part of these experiments may include variation between samples in bit-depth, spatial resolution, grid size, dye intensity, microscope channel definition or a variety of other parameters that are not critical to measuring quantities of interest but that do demand a software pipeline that supports programmatic configuration.

Open source tools similar to Cytokit include the <u>CODEX toolkit</u>, histoCAT [9], KNIME [10], ImageJ [11], Ilastik [12] and CellProfiler (CP) [13]. The CODEX toolkit is most similar but requires Windows as well as licensed deconvolution software, and it does not make use of common image processing libraries for segmentation or quantification so it is difficult to understand, debug, and extend. histoCAT is also very similar as it was built for analyzing multiplexed tissue images and the authors demonstrate how to use CellProfiler, Ilastik, and MATLAB to generate the data necessary for doing so. The primary drawback with this approach is that the operator needs to configure processing in a different graphical user interface (GUI) for each of the classification (Ilastik), segmentation (CellProfiler), and analysis (histoCAT) components of the pipeline. Cytokit aims to have a single configuration that can be applied to all downstream operations in an unattended manner, which makes it easier to run the same pipeline on similar data with only minor modifications (e.g. different channels). This comes at a loss of versatility because the provided segmentation method in Cytokit is not suitable for all imaging modalities, but custom segmentation routines can be integrated if they were developed externally. Out of the remaining tools, ImageJ, CellProfiler and Ilastik can all be utilized in the processing of multiplexed image data (as well as a multitude of other use cases Cytokit is not appropriate for) but none of them directly support this well as each would require a user to configure per-channel operations. This is a substantial burden when it means that some action needs to be taken in a GUI for each of potentially 20 or more channels, and it is particularly problematic when those channels change slightly in a new experiment. Programmatic configuration of such operations is more resilient to these changes and as an example of how Cytokit accomplishes this, CellProfiler is integrated as an option for image intensity quantification where the CP pipeline instance is generated and executed dynamically based on a static experiment configuration that can be altered without requiring the tedious and error-prone GUI interactions otherwise necessary with multiplexed image data. Lastly, KNIME is also similar to Cytokit in that it offers a way to implement control flow around native image processing operations as well as those provided by other projects. It is very likely that the same functionality in Cytokit could be captured by a comparable KNIME workflow, but Cytokit emphasizes support for offline, batch processing without a GUI-driven configuration -- a choice most likely to appeal only to bioinformaticians.

In summary, we present and validate a library providing programmatic configuration and GPU accelerated implementations of image processing algorithms for, but not exclusive to, analyzing high dimensional immunofluorescence data. This library is open source, requires nothing more than an existing <u>nvidia-docker</u> installation to run, is adherent to best practices such as continuous integration and unit testing, and includes a novel user interface alongside support for CellProfiler Analyst [14] to help navigate the cellular characteristics measured by this next generation of fluorescent imaging technology.

## Implementation

Cytokit consists of four major components, each of which will be discussed further in the following sections:

- **Processing** – Processing operations are applied to tiled images in parallel and in a predefined order to maximize ease of use as well as ensure that I/O for large image files is minimized by avoiding multiple reads and writes for the same images;
- **Configuration** – Beginning from a template configuration, image operations and extractions can be enabled/parameterized through “deltas” to concisely define variations on experimental results;
- **Extraction** – Cytometric single cell data can be exported as Flow Cytometry Standard (FCS) or comma-separated values (CSV) files and individual channels can be grouped into arbitrary subsets for extraction into ImageJ compatible Tagged ImageFile Format (TIFF) hyperstacks for custom analysis;
- **Visualization** – The Cytokit Explorer UI allows for Individual cells to be isolated based on phenotype and visualized within the entire field of view for an experiment, individual fields of view, or as single cells.

### Processing

The imaging processing pipeline in Cytokit, based largely on the original <u>CODEX toolkit</u>, is designed to support datasets with the following dimensions:

- Tile – A single field of view
- X/Y/Z – 3D images captured at each tile location
- Channel – Images for different expression markers
- Cycle – Groups of channels (usually 3 or 4) captured between dye exchange cycles
- Region – Groups of tiles often collected as square grids

This seven dimensional structure, illustrated in **Figure 1**, is common in multiplexed imaging but dimensions of length 1 are also supported in nearly all cases. In other words, the smallest Cytokit-compatible experiment would consist of a single, grayscale 2D image with a height and width exceeding some minimum value (currently 88 pixels). All of the imaging processing steps applied to these datasets can be enabled or disabled based on the experiment configuration and are executed in a predetermined order. The individual steps themselves are implemented using existing deep learning models or are ported from other Java or Python libraries to run as GPU-accelerated TensorFlow [15] computational graphs. There are some exceptions to this such as image I/O and cell segmentation post-processing, but all of these are discussed below.

**Figure 1:**
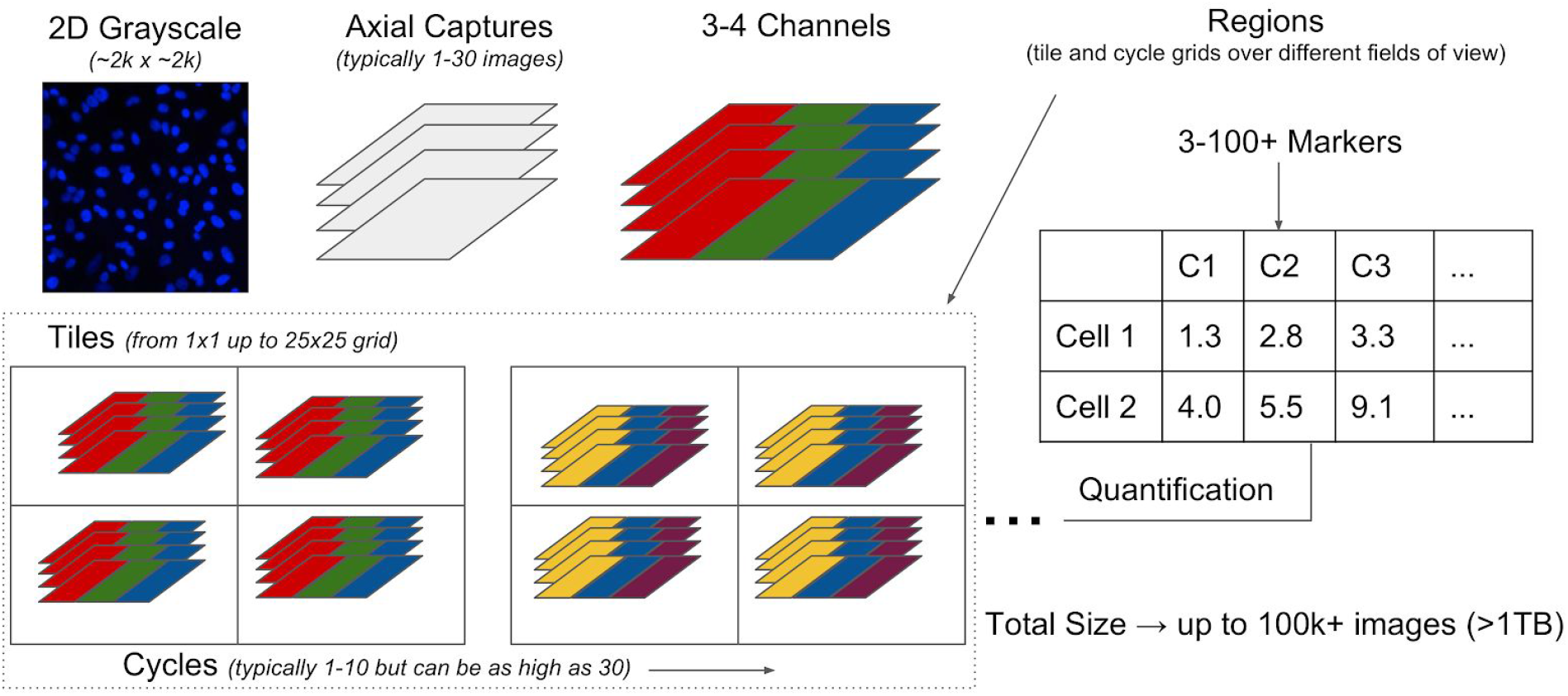
Seven dimensional multiplexed experiment structure illustrating how 3D single channel images are grouped as cycles, captured as tiles on a grid, and then potentially repeated over multiple fields of view (as “regions”) before being quantified as two dimensional single-cell information.

#### Operations

As illustrated in **Figure 2**, the first step in the Cytokit pipeline involves loading images into memory in a way that is decoupled from all further processing steps. This is done using a separate thread, outside of any TensorFlow graphs, to assemble 2D grayscale images as 5 dimensional tile arrays that are then loaded onto a queue of configurable size (typically 1). This ensures that as processing for a single tile completes, there is no delay introduced by disk I/O before beginning processing on the next tile. This is an important feature for maximizing GPU utilization and reduces overall processing time substantially when the time necessary to process a single tile is not drastically larger than the time necessary to load it from disk. The image files themselves are assumed to be 8 or 16-bit 2D grayscale images with file names containing region, tile, cycle, z-plane, and channel index numbers, in a configurable format, so that the array structure matches that of the experiment.

**Figure 2:**
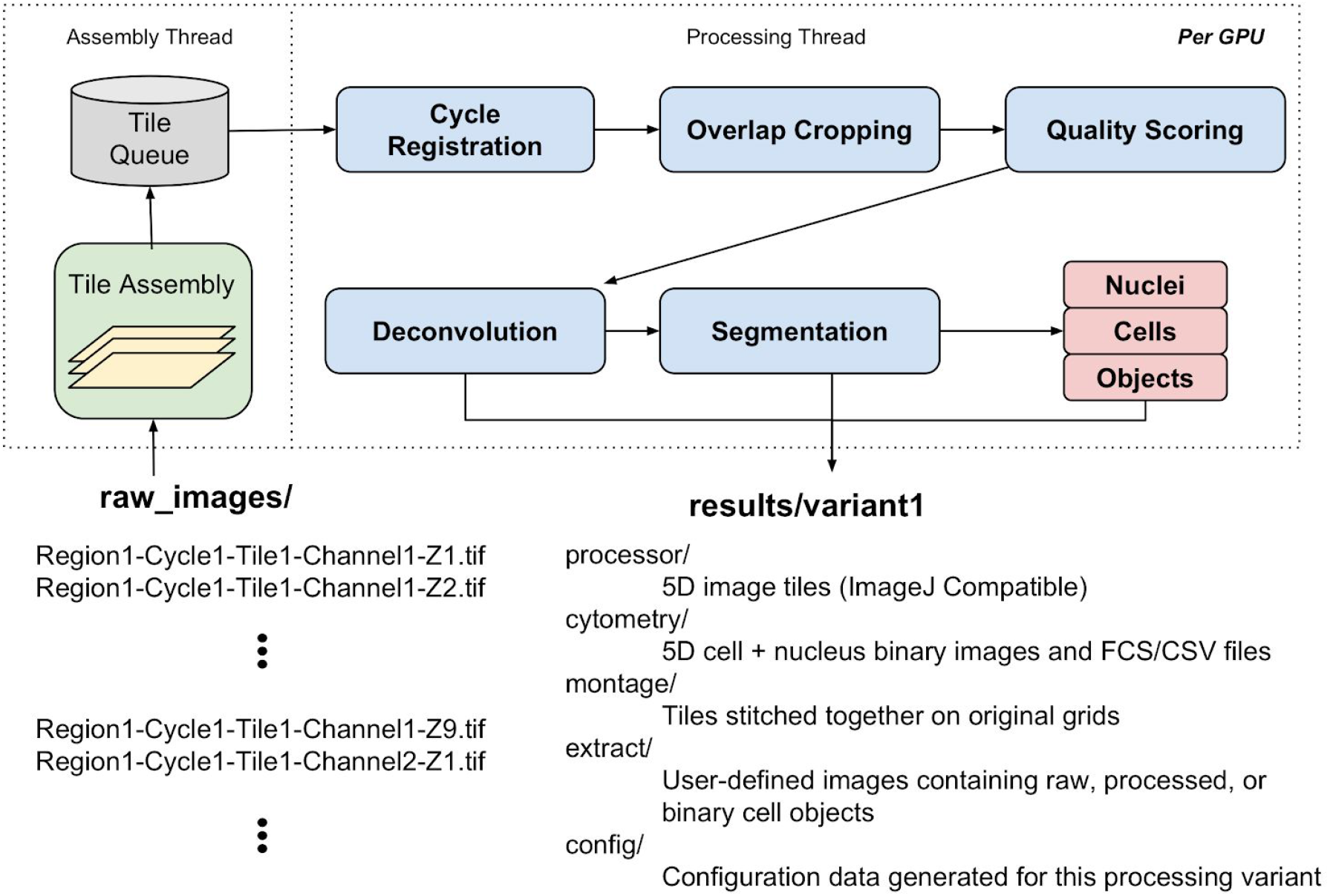
Processing pipeline overview with operations in original CODEX project replaced by GPU-accelerated equivalents, decoupled from tile assembly, and modified to support labeled object extraction as well as ad-hoc image stacks/montages

The remaining image manipulation operations include:

##### Cycle Registration

This operation serves as a way to align reference images across cycles and then apply the inferred translation to all other non-reference images. As is necessary in multiplexed imaging, a reference channel usually containing a nuclear stain must be collected in each cycle so that the sample drift occurring in the time elapsed between fluorophore exchanges can be compensated for in downstream analysis. In Cytokit, this operation is implemented as a port of the cross correlation algorithm [16] in scikit-image [17] `register_translation` to TensorFlow.

##### Image Quality Assessment

This is provided by the <u>Microscope Image Quality</u> [18] project, a TensorFlow classifier that allows for images to be scored based on their quality, and is used in Cytokit to select individual 2D images for visualization and/or quantification.

##### Deconvolution

3D image deconvolution in Cytokit is provided by the <u>Flowdec</u> project, which is a direct port of the Richardson Lucy algorithm in the DeconvolutionLab2 [19] library to TensorFlow. Cytokit also includes support for automatically generating Point Spread Functions based on an experiment configuration through a fast Gibson-Lanni kernel approximation method [20].

##### Segmentation

Identification of cell nuclei is performed using the deep learning model [21] featured in CellProfiler. The semantic segmentation provided by this model produces a 3 class prediction for background, nucleus interior, and nucleus boundary pixel classification. A nuclei object image is generated from these predictions as a single pixel dilation of binary objects resulting from the selection of the pixels that have the highest probability of belonging to the nucleus interior class, after removing nuclei below a configuration size threshold. The nuclei objects are then used to identify entire cell objects based on either a fixed radius outside the nucleus or, if available, a membrane stain channel used to create a threshold image serving as a watershed mask or as both a mask and a “height” image for propagation segmentation [22] (to trade off distances between nuclei with membrane image intensity in the formation of cell boundaries). Any intermediate image processing subsequent to segmentation is implemented using scikit-image, SciPy [23], and OpenCV [24]. Quantification of expression images over the area of the segmented objects is also performed using modules in CellProfiler. This operation will produce files compatible with either Cytokit Explorer or CellProfiler Analyst.

#### Configuration

Experimental variation in high throughput microscopy imaging often requires processing pipelines that are very tunable. While the intention in Cytokit is to employ algorithms that require this as little as possible, some configurability is unavoidable. The approach taken to address this problem involves two core capabilities -- iteration and evaluation. Much like any general hyperparameter optimization process, Cytokit attempts to make defining iterations and evaluating them as simple as possible so that tuning individual operations for individual images is much less common than being able to view the effects of parameter settings across an entire experiment. For example, a common process in our lab is to generate a template configuration that is then modified incrementally to produce several variants of an experiment for analysis. A simple version of this process is shown in **Figure 3** and demonstrates how the volume of data incorporated may be increased as appropriate parameter settings become clearer.

**Figure 3:**
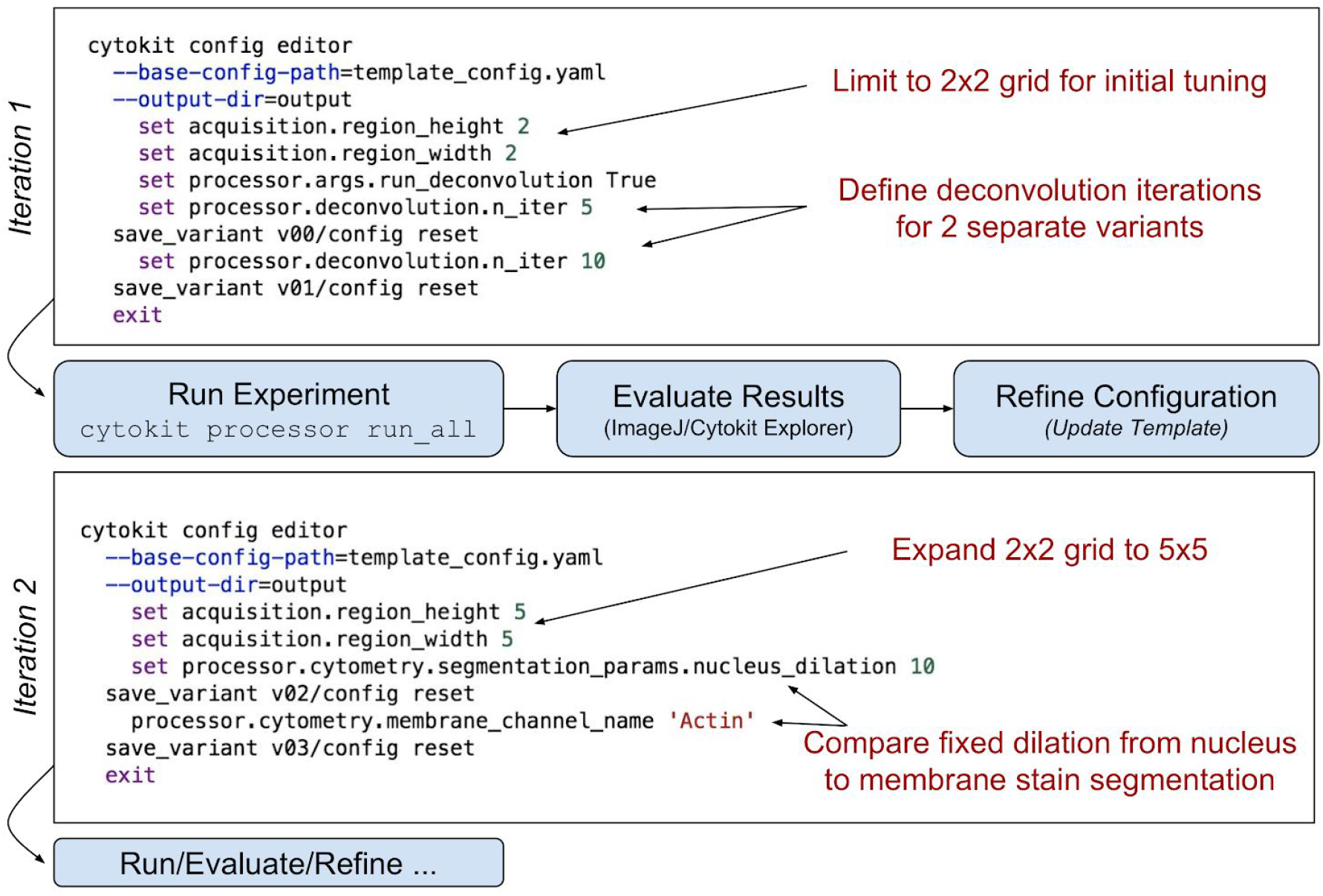
Example iterative pipeline optimization process with CLI commands used to continually refine and expand the scope of experiment processing for large raw image datasets

Template configurations, which may also simply be the sole configuration for an experiment, need to at least include the dimensions of the experiment (grid size, number of Z planes, image overlap, etc.), microscope parameters, and channel names. They may also contain definitions for any commands to be run as a way to ensure that all processing for an experiment is defined in one place. Practically speaking, this means that arguments to a command line interface (CLI) controlling processing, data extraction, or analysis may be specified in the experiment configuration or overridden at the command line. This often fosters reproducibility since it eliminates the need to keep track of how a pipeline was invoked while still making it possible to introduce small changes in behavior. A good example of this is defining channel subsets to extract for visualization as this can be done completely ad-hoc at a command line or defined in the template configuration if the extraction is commonly useful.

### Extraction

Extracting data from high dimensional experiments can be challenging, particularly when slices of interest across those dimensions span image volumes that are too large for other visualization and analysis software. Extraction utilities in Cytokit make it possible to mix raw image data with processed image results as well labeled object data (for cells/nuclei) and can be parameterized to operate on subsets of image grids or fields of view, lists of specific channel names, or z-plane subsets. Those extracted images can then also be stitched together such as in **Figure 4** where a 49 tile (7×7 grid) montage of 1008×1344 images was generated as single TIFF file (cropped to 8192×8192 pixels) containing marker expression levels and cell/nucleus boundary data.

**Figure 4:**
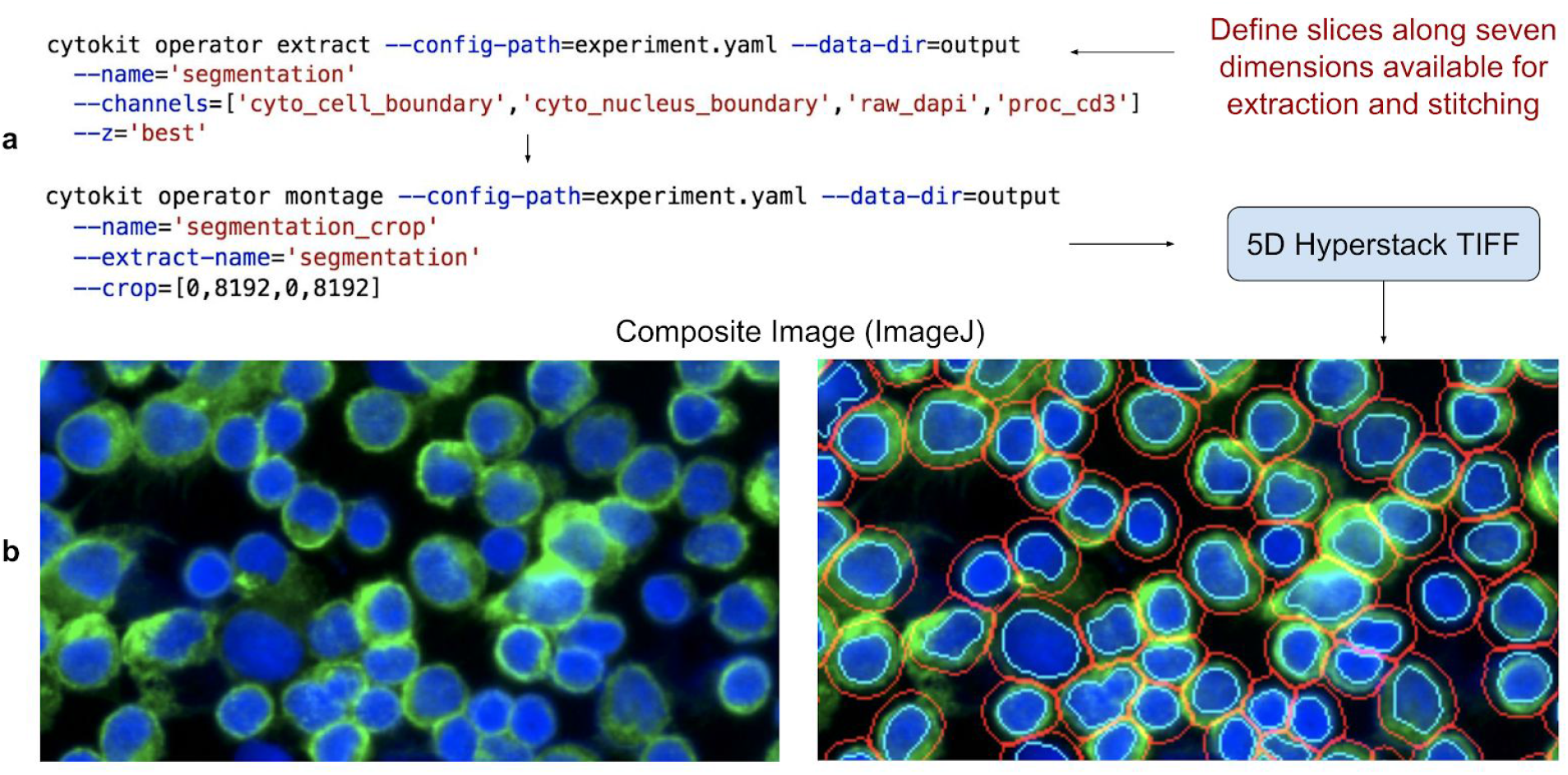
Example extraction and montage CLI commands. **(a)** CLI commands define slices as names (where relevant) or as lists and ranges of indexes to extract raw, processed, or object image data. **(b)** Resulting TIFF files are ImageJ compatible for blending and visualization, like the example shown with human T cells labeled as CD3 (green), DAPI (blue), nuclei boundaries (cyan), and cell boundaries (red).

In addition to image extraction utilities, Cytokit also offers single cell data as FCS or CSV files containing cell identifiers linking back to object images, location coordinates within the experiment, morphological properties (diameter, size, circularity, etc.), mean expression levels across the entire cell or within the nucleus alone, and graphical properties like identifiers of adjacent cells, number of adjacent cells, and size of contact boundary. The same information (and more) can also be exported using CellProfiler directly to generate a SQLite database compatible with CellProfiler Analyst.

### Visualization

One of the primary challenges in in-situ image cytometry is developing an understanding of the relationship between phenotypic and spatial properties of cells. Multiplexed imaging further complicates this process as the quantity of images produced both increases the likelihood that illumination artifacts exist along the spatial dimensions and often makes finding them manually infeasible. To assist in interrogating these relationships as well as build phenotypic profiles for cell populations, an interactive Dash (by Plot.ly) application is provided that allows for gates applied to cytometric data to be projected onto images within an experiment. Shown in **Figure 5**, this application can be used to visualize individual cell/nucleus segmentations, highlighted as SVG overlays, within the context of a stitched grid view of an experiment as well as a single field of view. Additionally, single cell images can be extracted from an entire experiment with blended overlays of various expression channels (having user defined contrasts) as a means of ensuring that the characteristics of cells assumed to exist in any one population gated purely based on numerical information match expectations. Custom metrics can also be computed and attached for visualization in this interface using any of the fields mentioned in the “Extraction” section above.

**Figure 5:**
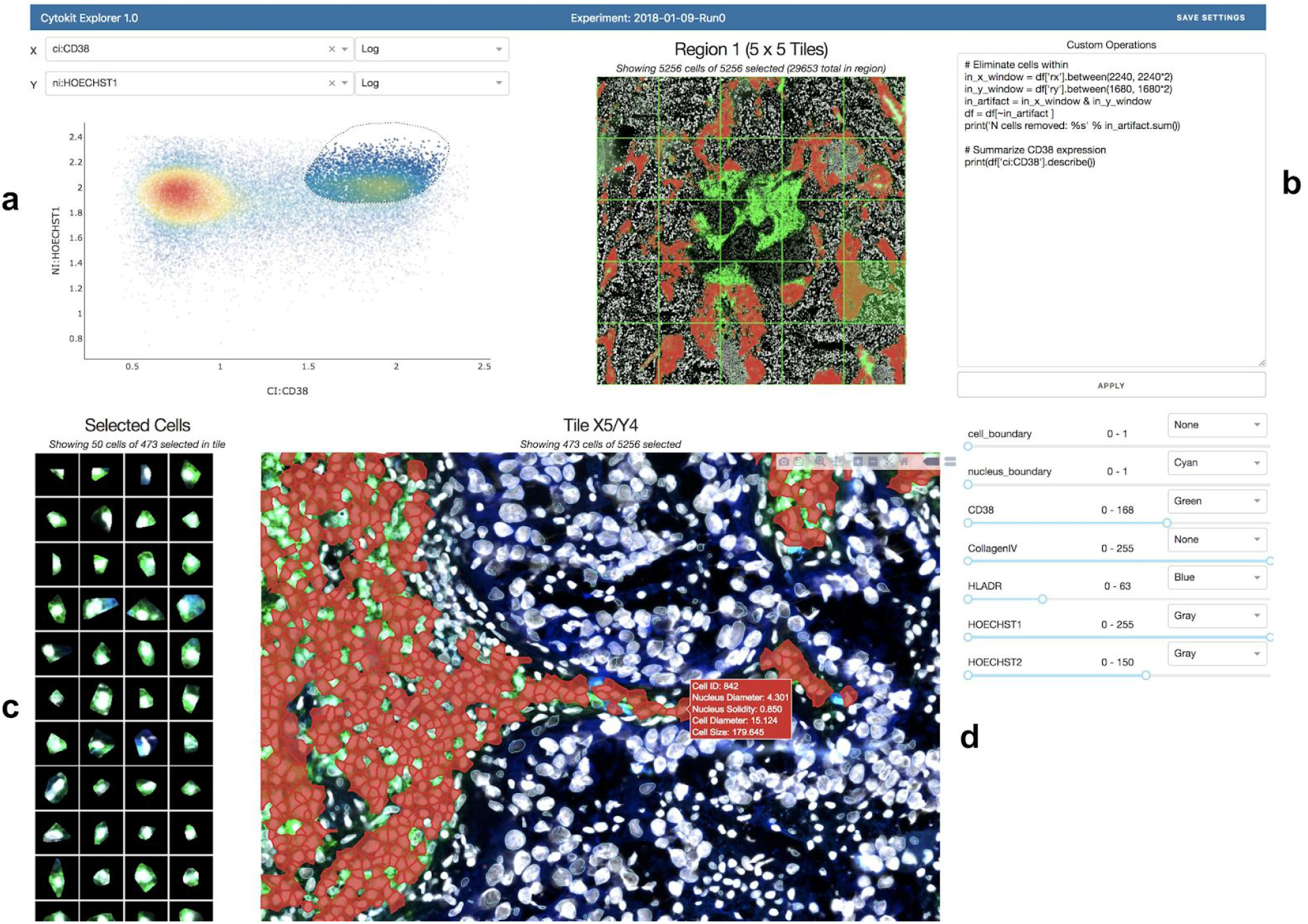
Cytokit Explorer screenshot (see <u>screencast</u> for animated version) showing a CODEX sample imaged at 20x. **(a)** 1D or 2D plots of expression, morphological, or graphical cell features support box and free-hand gating. **(b)** Custom filtering, to ignore a central photobleached region of cells in this case, or summarizations are applied immediately. **(c)** Single cell images match current gate and selected channel display settings and can also be buffered onto the page as tiles are selected, or across the entire image grid (not shown). **(d)** Gated cell population projected onto selected tile image with current display settings

## Results

### Cellular Marker Profiling

To demonstrate the extraction and analysis features of Cytokit as well as validate the underlying image processing libraries, a series of traditional immunofluorescence experiments were first conducted on human primary T cells. The first of these, shown in **Figures 6** and **7**, comprised of primary human T cell samples stained with CODEX oligonucleotide-conjugated antibodies against human CD3, CD4, and CD8 (and a separate HOECHST stain). These slides were then imaged at 20X on a 1.9mm × 2.5mm grid of 25 images over an axial depth of 12.5 micrometers. The resulting 1008×1344×25 (height × width × depth) image volumes were then deconvolved, segmented, and quantified before being gated using Cytokit Explorer to isolate helper (CD4 positive) and cytotoxic (CD8 positive) subpopulations.

**Figure 6:**
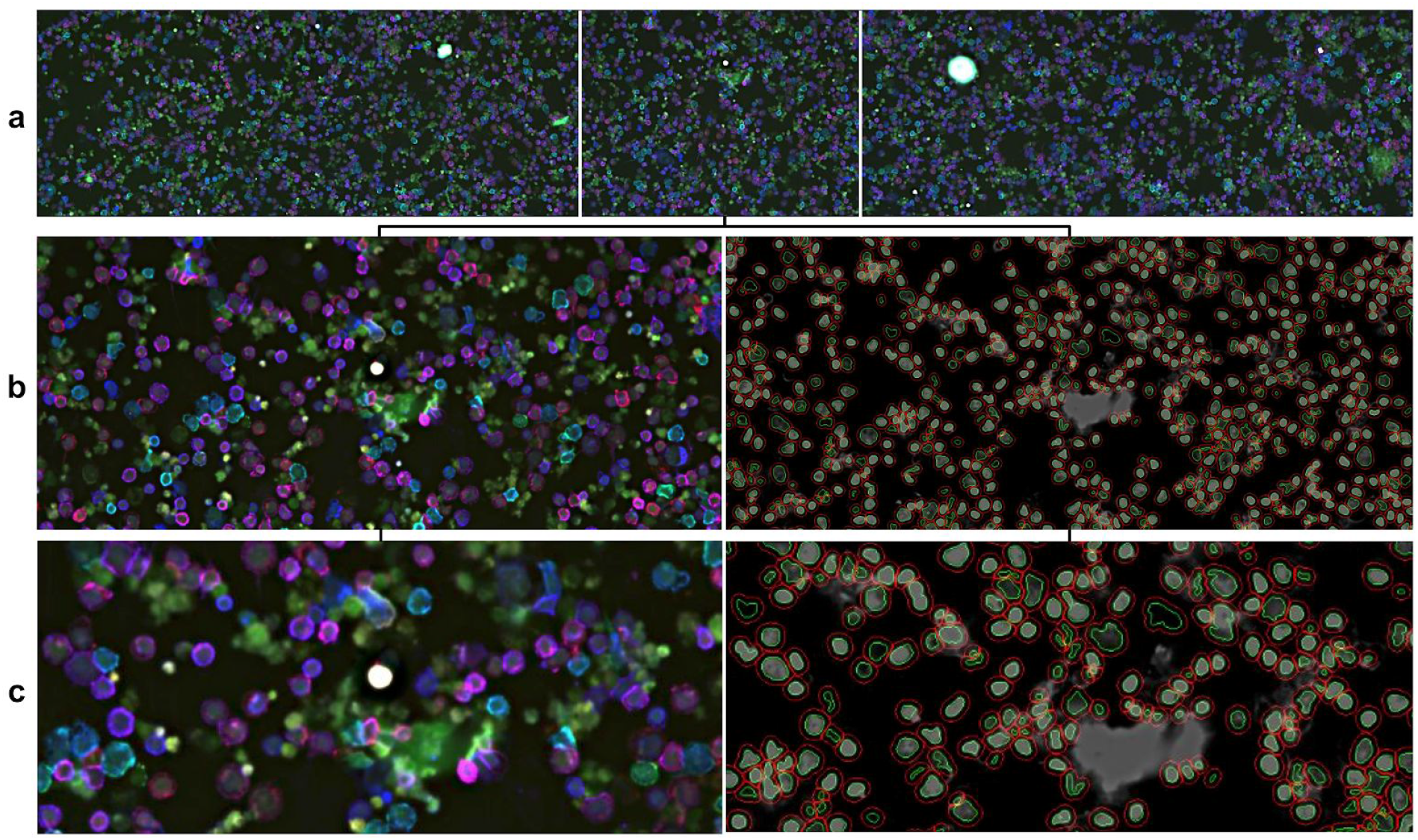
T cell CD3 (blue), CD4 (red), and CD8 (green) intensity. **(a)** First row of 5 images in 5×5 experiment grid. **(b)** Single tile image with corresponding cell and nucleus segmentation, where cells are defined as a fixed radius away from the nucleus in the absence of a plasma membrane stain. **(c)** Center zoom on (b) showing co-expression of CD3 and CD4 (magenta) and CD3 and CD8 (cyan) as well as debris in DAPI channel.

**Figure 7:**
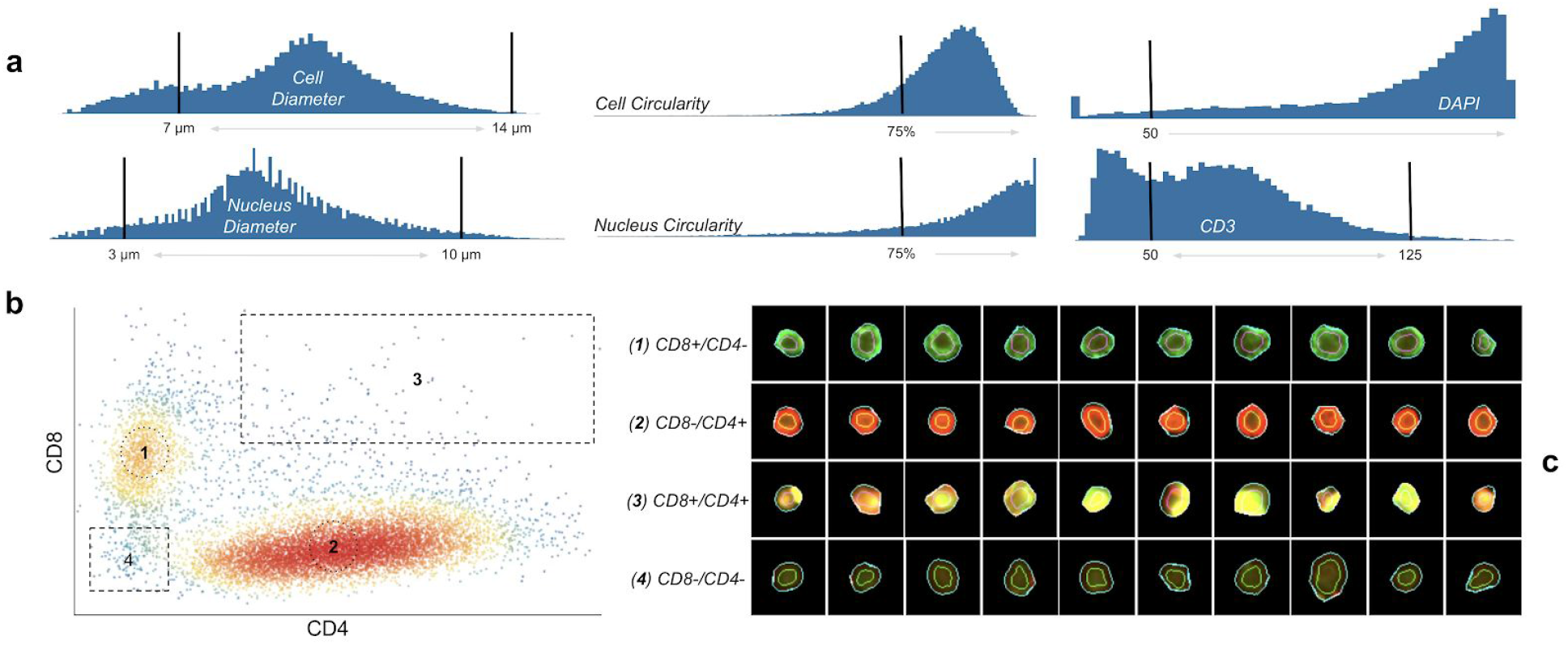
T cell gating workflow as annotated Explorer screenshots. **(a)** Morphological and intensity gates applied to isolate CD3+ cells. **(b)** Cytotoxic and helper cell subpopulations. **(c)** Individual cell images matched to subpopulations in (b).

While these CD4 and CD8 positive (i.e. CD4+ and CD8+) populations were easily resolved in this experiment (**Figure 7**), we found that dissociated cell samples like this are difficult to prepare without non-trivial amounts of debris and diffuse nuclear staining, usually as a result of lysed cells that did not survive the centrifuge. This is visible in **Figure 6 (c)** where a minority of the nuclei segmentations resulting from the CellProfiler U-Net are fixed around roughly circular areas of greater DAPI intensity, albeit at low contrast. This invariance to contrast is generally very desirable, but it also demonstrates the importance of curation in image cytometry as artifacts like this can easily go undetected without a way to relate variations in inferred morphological or expression profiles for cells back to raw images.

A further investigation of the ability of this method to isolate CD4+CD8- and CD4-CD8+ cell populations was also conducted on experimental replicates and validated against flow cytometry based surface marker profiling. Shown in **Figure 8**, population proportions matched closely and verified that dissociated cells quantified in this way can produce results comparable to other methods; however, tools like Explorer and OpenCyto [25] were necessary to reach this degree of parity due to over-saturated image tiles (detected in Explorer) and intensity calibration differences resulting in substantial movement in the modes of the CD4/CD8 populations across replicates and donors. This latter issue was compensated for, in a downstream analysis, through the use of the t-distributed mixture models provided in OpenCyto (via flowClust [26]) that can capture translated distributions regardless of the intensity scale unique to each imaging dataset.

**Figure 8:**
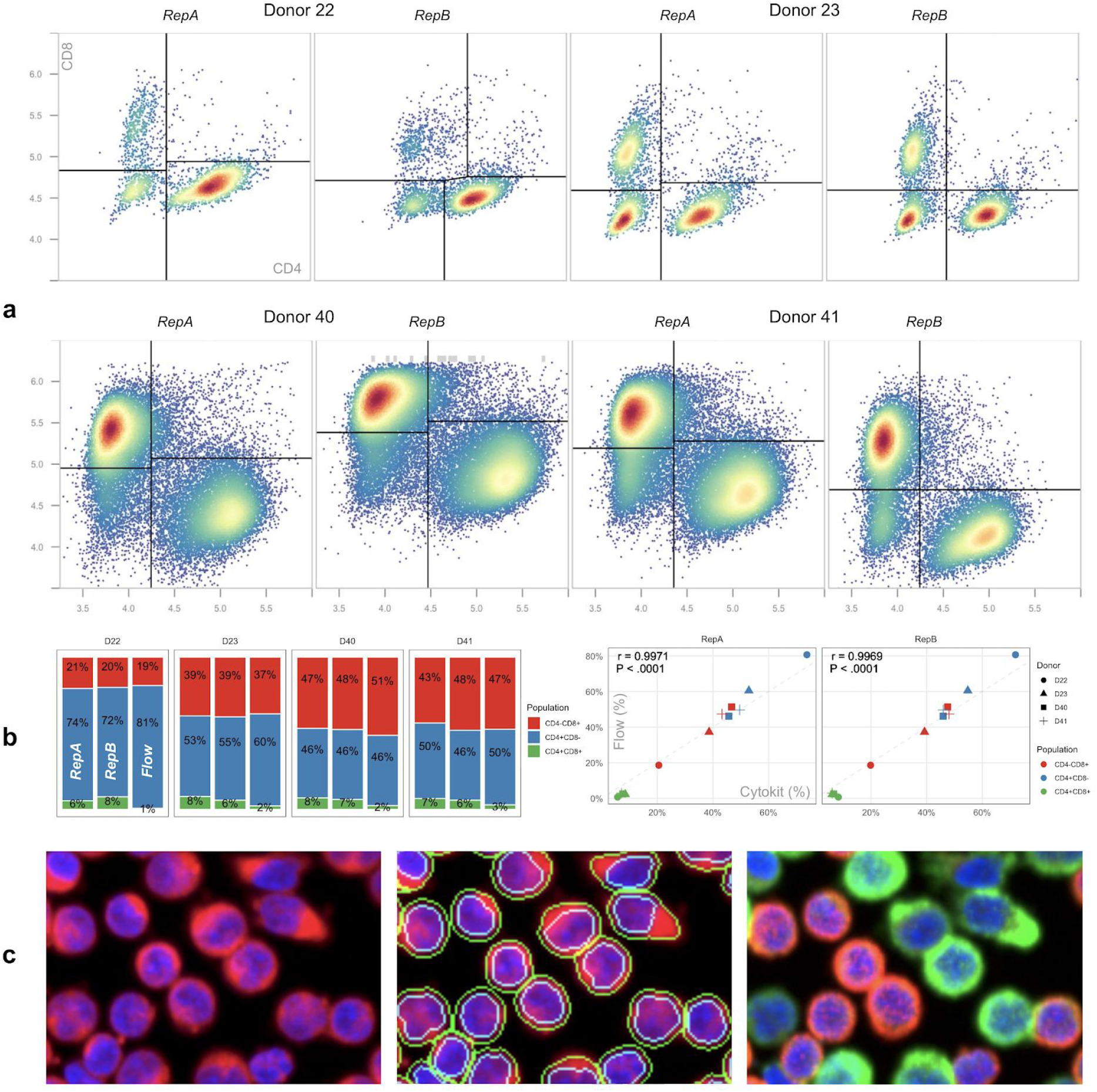
T cell population recovery comparison (<u>notebook</u>). **(a)** CD4/CD8 gating results, as determined by automatic gating functions in OpenCyto [25], over two imaging replicates for each of 4 donors. While all images were collected over a field of view of the same size, samples for donors 40 and 41 were prepared at 3x higher cell concentrations to demonstrate that segmentation and intensity measurements are robust to greater image object densities. **(b)** Cell population size for both replicates compared to a single flow cytometry measurement for each donor as well as Pearson correlation demonstrating strong agreement between the two (r > 0.99, P < 0.0001, two-tailed t-test). **(c)** Cell images from donor 41 showing (from left to right): DAPI (blue) and PHA (red) stain, DAPI and PHA with cell and nuclei segmentations, and DAPI with CD4 (red) as well as CD8 (green). See supplementary file **Figure S1** for a comparison of these results to those from the same workflow without much of the gating used here to remove invalid cells.

### Cell Size Estimation

A second validation was also conducted to attempt recovery of known morphology differences between unstimulated and activated T cell samples. Samples were stained with DAPI and Phalloidin-Fluor 594 (PHA), the latter of which identifies cytoskeletal actin filaments and is used here to define cell boundaries. The Cytokit pipeline for processing both samples was configured to blur PHA images before applying a threshold to create a binary image used as a watershed segmentation mask, with nuclei as seeds. As in the previous experiments, Explorer was used to find appropriate gates for singlet cell populations before attempting to compare cell diameter distributions (**Figure 9**).

**Figure 9:**
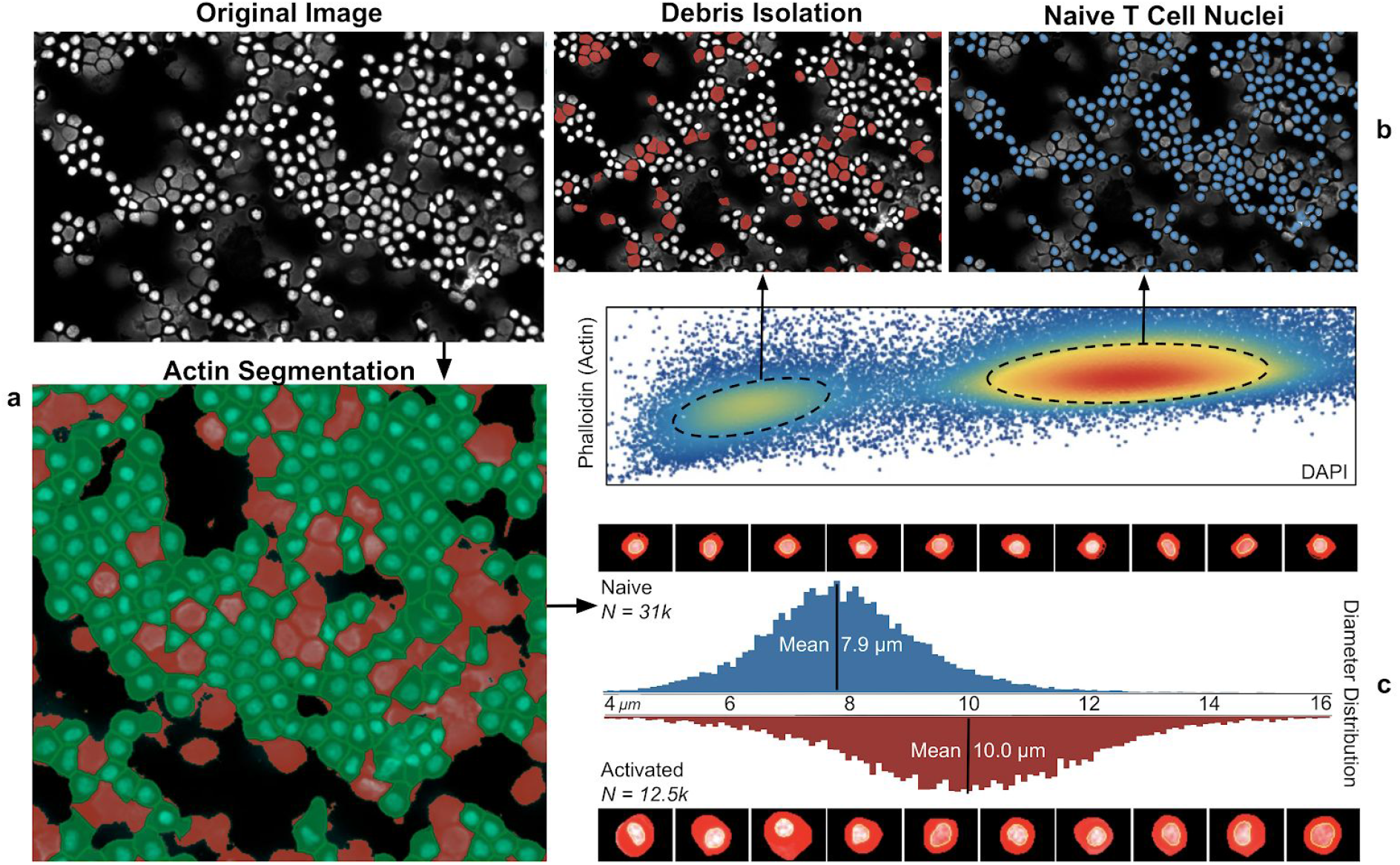
Cell diameter recovery workflow as annotated Explorer screenshots. **(a)** Unstimulated/naive cells with DAPI stain (gray), binary PHA image (red), and resulting cell segmentation (green). **(b)** Projection of modes in phalloidin/DAPI distribution to segmented cells in original image. **(c)** Diameter distribution comparison for unstimulated and activated samples along with corresponding single cell images.

Much like the previous section demonstrating estimation of CD4+/CD8+ T cell population sizes, obtaining accurate cell size distributions was found to be difficult without first establishing proper filters to eliminate debris, lysed cells, and poorly segmented nuclei. Similar procedures for assays based on dissociated cells, e.g. Flow and Mass Cytometry, are far more formalized yet satisfactory results can still be obtained with image cytometry as long as the relationships between spatial and expression dimensions can be properly characterized. As this characterization becomes more difficult across different experimental replicates and modalities, a series of similar experiments were also carried out to better define these difficulties on a larger scale. **Figure 10** demonstrates results from these experiments where a workflow based on automatic gating functions, with resulting filters validated against individual cell images in Explorer, was used to extract cell size distributions for different cell types, levels of magnification, and replicates for comparison to similar results from a dedicated cell counting device.

**Figure 10:**
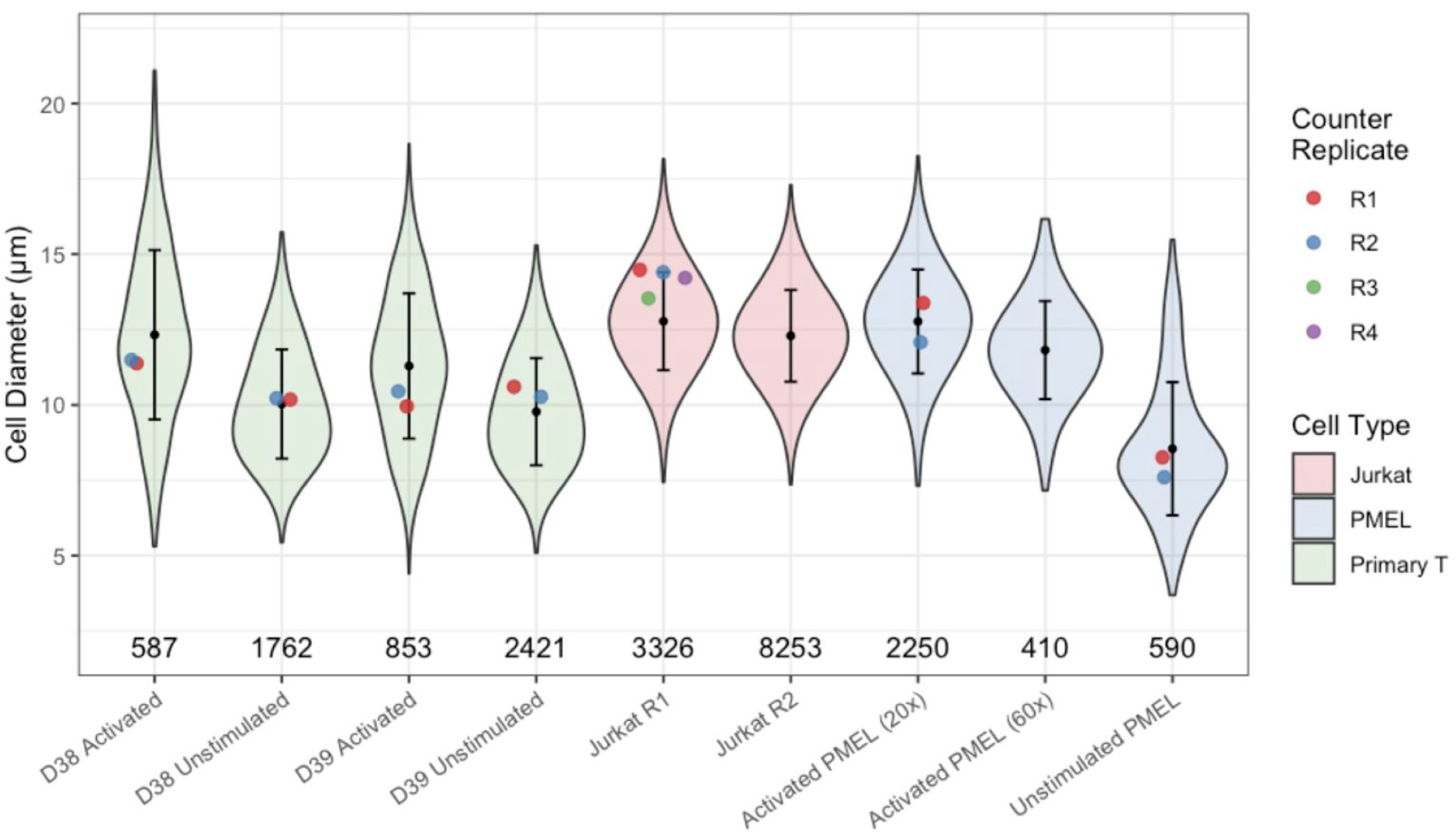
T cell size recovery comparison (<u>notebook</u>) showing inferred cell diameter distributions (violin with bars indicating +/− 1 s.d.) vs point estimates of mean diameter from Thermo Fisher cell counter (dot) as well as filtered single cell population sizes (counts); All images taken at 20x magnification except where indicated otherwise. See supplementary file **Figure S2** for a comparison of these results to those from the same workflow without much of the gating used here to remove invalid cells.

### CODEX

Representative preparations of image datasets resulting from CODEX protocol applications were also carried out to characterize performance at a larger scale. The first of these included an analysis of the <u>reference dataset</u> shared by the CODEX authors which, in its totality, contains ~700k cells spread across 9 individual tissue preparations and ~1.1TB of microscope images. These images, of normal and autoimmune murine spleens, were prepared as separate replicates of tissue slides with a single replicate being subjected to 18 cycles of imaging (on a 9×7 grid) to capture 54 expression signals. Cytokit was applied to a single replicate containing 51,030 images (129GB) to demonstrate that cycle alignment, deconvolution, image quality assessment, and segmentation can be executed in ~80 minutes on one (2× Nvidia GTX 1080 GPU) workstation. Segmentation results*^1^ were visually and quantifiably similar to those in the original study (**Figure 11**).

**Figure 11:**
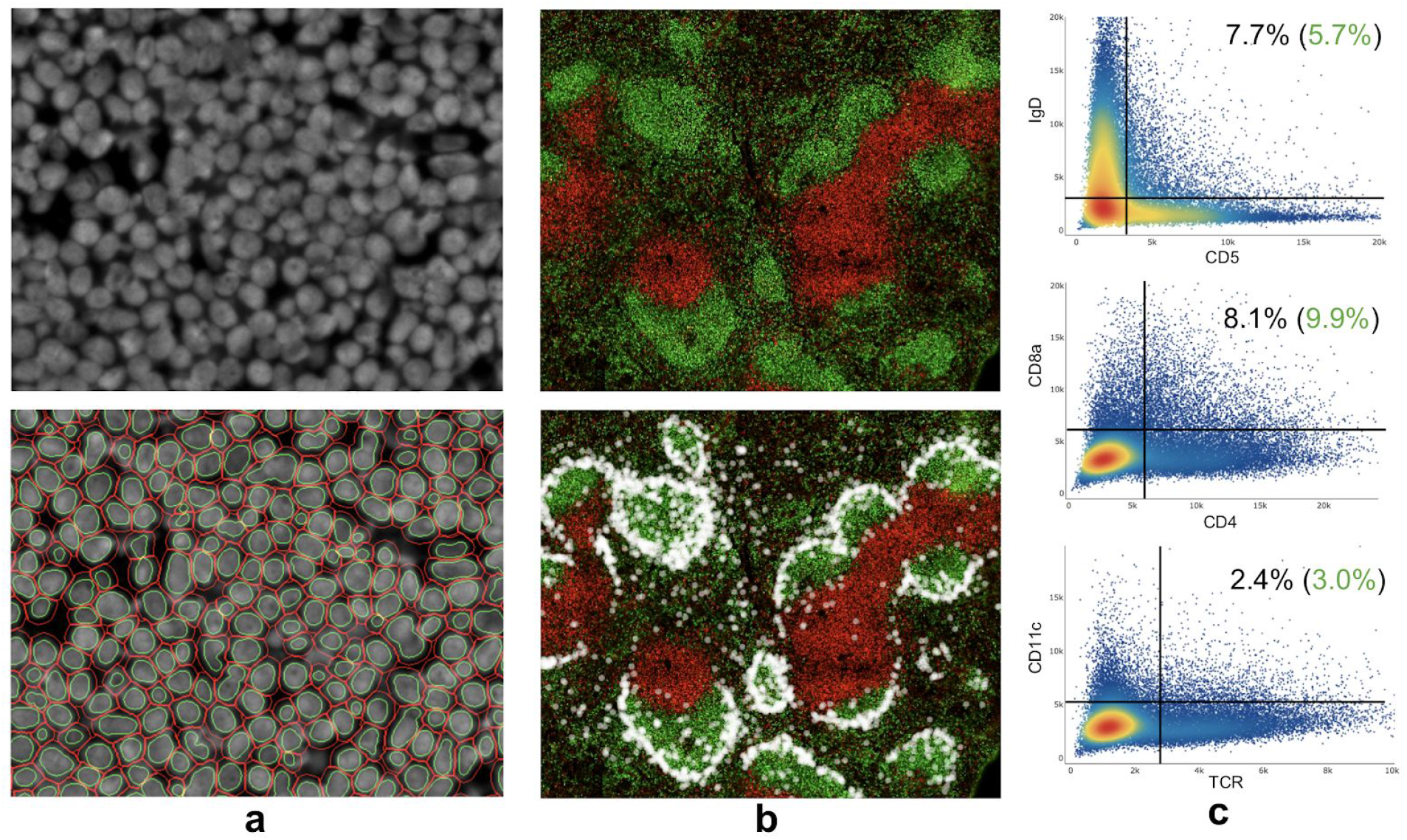
BALB/c spleen (slide 1) CODEX image segmentation/quantification results. **(a)** ~200 cells with DRAQ5 nuclear stain (above) and corresponding cell/nucleus segmentation (below). **(b)** ~71k cells in stitched, downscaled 9072×9408 pixel image showing IgD (green) and CD90 (red) expression as well as location of CD169+ marginal zone macrophages as white dots. **(c)** Double positive cell population rates post cleanup-gating with expected percentages in green

## Conclusion

Multiplexed, in-situ image cytometry offers many potential advantages over traditional single cell assays, but the volume of image data produced by these procedures makes processing and analyzing them difficult. Cytokit is an open source Python package that attempts to help overcome these issues by providing GPU-accelerated implementations of cycle registration and image deconvolution as well as an interactive user interface for characterizing the phenotypic features of cells alongside their spatial distribution. We find that understanding modalities in expression distributions as projections onto original images and the ability to iteratively optimize pipelines operating on large datasets are both important capabilities needed to achieve good results, even for the relatively simple tasks presented here such as size estimation and cellular marker profiling. We also find that the need for this software arises largely out of the inability of existing software packages to support experiments with large numbers of imaging channels well. It is likely that as CellProfiler continues to mature, it will eventually support a Python API that would make it possible to provide many of the same features with CP modules alone. Until then, Cytokit is an efficient, flexible pipeline suitable for bioinformaticians that need to process and analyze multiplexed imaging data.

## Supplementary Figures

**Figure S01:**
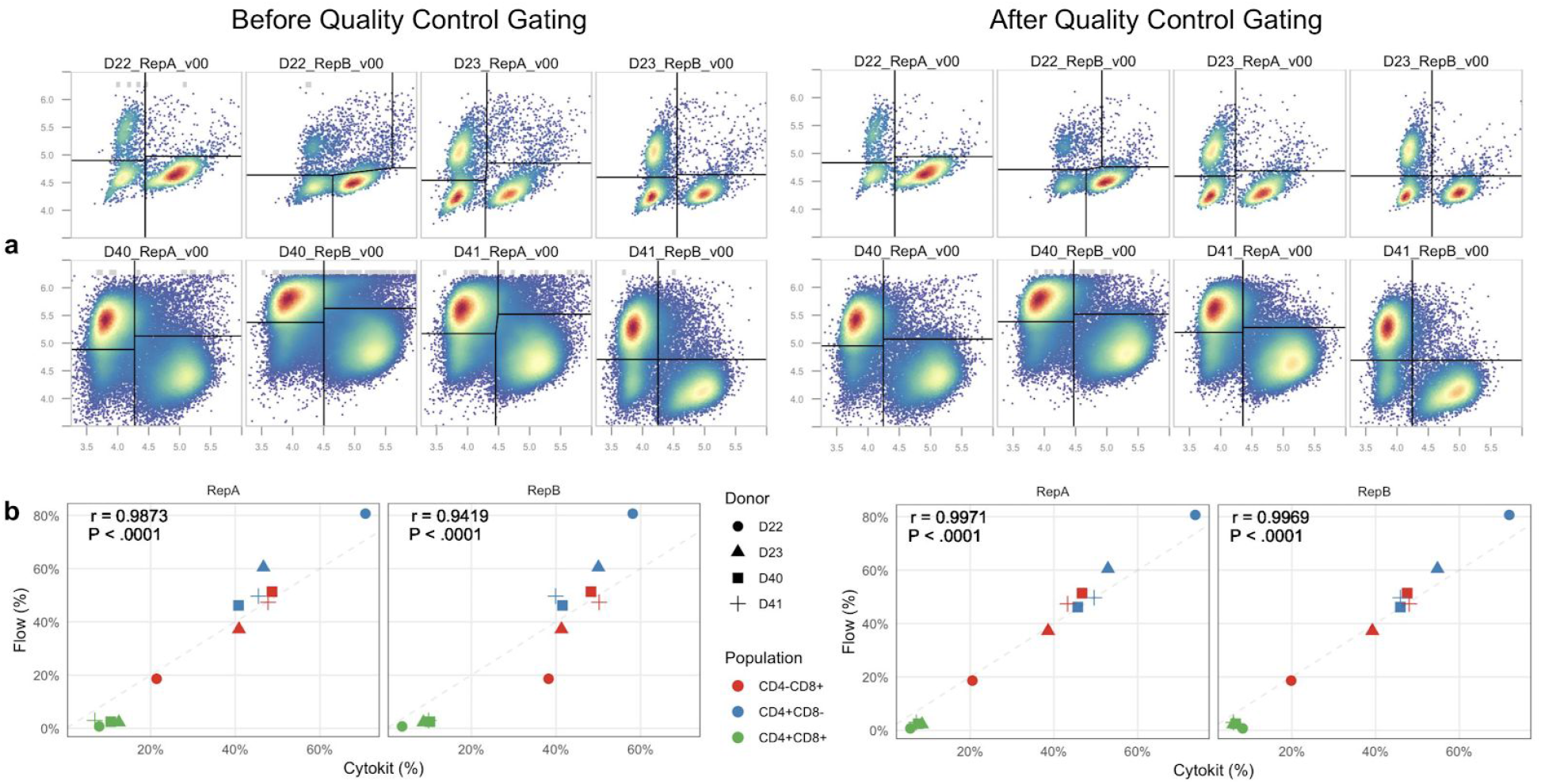
Supplement to **Figure 8** demonstrating improvement in quantification after the inclusion of gates to remove debris and other segmentation artifacts. **a)** CD4/CD8 cell populations for all 4 donors with both before and after groups including a terminal gate in the workflow to detect modes for each population, but all other gates left out in the “Before Quality Control Gating” example. **b)** Cell population size for both replicates compared to flow cytometry measurements with improvement in correlation across populations after application of quality control gating (pearson correlation shown with significance from two-tailed t-test).

**Figure S02:**
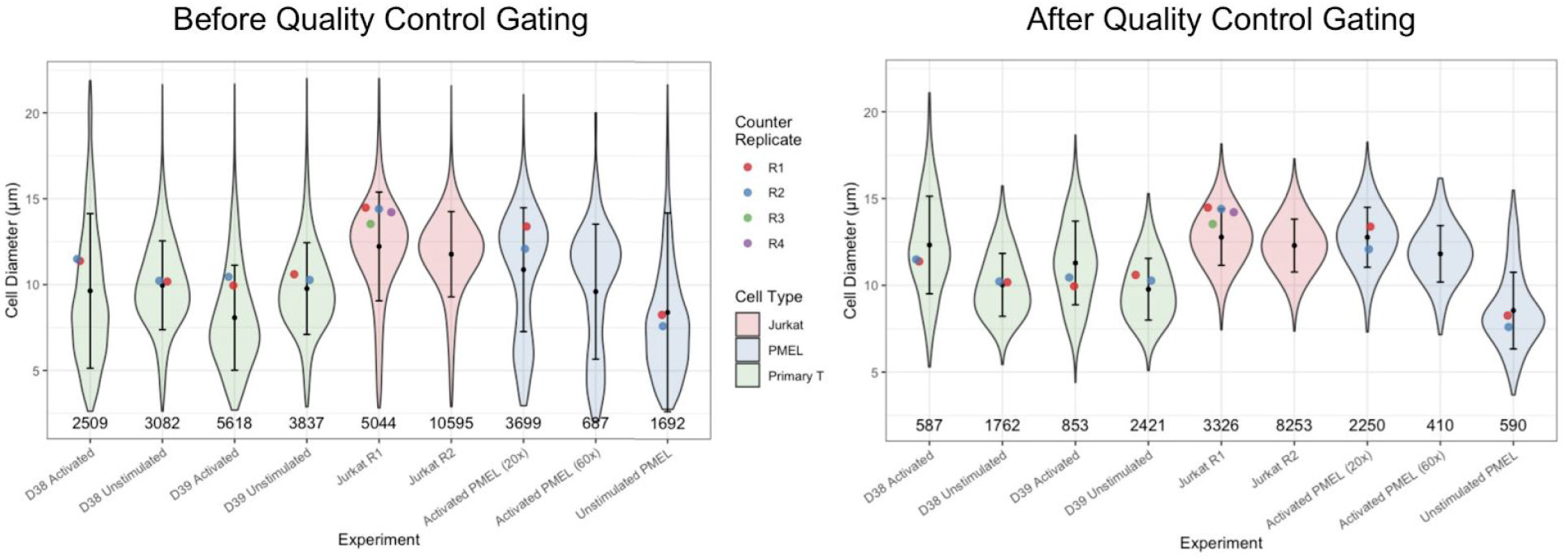
Supplement to **Figure 10** demonstrating improvement in quantification after the inclusion of gates to remove debris and other segmentation artifacts. Cell counts from the two workflows are shown below each diameter distribution (violin with bars indicating +/− 1 s.d.) and corresponding Thermo Fisher cell counter diameter measurements are shown as colored dots.

## Methods

For all human primary T cell imaging datasets generated in this study, cells were isolated, cultured, activated, and prepared for immunofluorescence microscopy as previously described [27]. PBMCs were isolated from healthy human donors (purchased from Plasma Consultants LLC, Monroe Township, NJ) by Ficoll centrifugation (Lymphocyte separation medium; Corning, Corning, NY). T cells were isolated using Dynabeads Untouched Human T Cells Kit using manufacturer’s protocols (Thermo Fisher, Waltham, MA). Isolated T cells were kept in T cell media: RPMI with L-glutamine (Corning), 10% fetal bovine serum (Atlas Biologicals, Fort Collins, CO), 50 uM 2-mercaptoethanol (EMD Millipore), 25 mM HEPES (HyClone, GE Healthcare, Chicago, IL), 1% Penicillin-Streptomycin (Thermo Fisher), 1X sodium pyruvate (HyClone, GE Healthcare, Chicago, IL), and 1X non-essential amino acids (HyClone, GE Healthcare). T cells were activated for 2 days with anti-CD3/CD28 magnetic dynabeads (Thermo Fisher) at a beads to cells concentration of 1:1, with supplement of 200 IU/ml of IL-2 (NCI preclinical repository) and then mounted onto a glass slide with the Prolong anti-fade mounting reagent. For samples stained with only DAPI and phalloidin, T cells from cultures were washed and re-suspended in PBS, fixed in 4% formaldehyde solution for 30 minutes, washed and stained with the phalloidin dye for 30 minutes within BD CytoPerm solution, and re-suspended in PBS before cytospinning. T cells from pmel-1 mouse were activated by adding 1 uM of gp100 peptide to freshly isolated splenocytes and culturing for three days.

### Protocol Details

Culture media: DOI:10.17504/protocols.io.qu5dwy6

PBMC isolation from buffy coat: DOI:10.17504/protocols.io.qu2dwye

FL staining: https://www.protocols.io/view/preparing-primary-t-cells-for-fluorescence-microsc-vede3a6

## Availability and requirements

**Project name**: Cytokit

**Project home page**: https://github.com/hammerlab/cytokit

**Operating system(s)**: Linux, Mac OS X

**Programming language**: Python

**Other requirements**: Python 3.5 or higher

**License**: Apache 2.0

**Any restrictions to use by non-academics**: None beyond license terms

## Abbreviations

CODEX: Co-detection by indexing
GUI: Graphical User Interface
PHA: Phalloidin-Fluor 594
CP: CellProfiler
DEI: DNA Exchange Imaging
TIFF: Tagged ImageFile Format
FCS: Flow Cytometry Standard
CLI: Command Line Interface
I/O: Input / Output

## Declarations

### Ethics approval and consent to participate

Not applicable.

### Consent for publication

Not applicable.

### Availability of data and materials

The datasets generated for this study are available at https://console.cloud.google.com/storage/browser/cytokit/datasets and the related analysis as well as configurations necessary for reproduction can be found at https://github.com/hammerlab/cytokit. CODEX data used for comparison to results in the original publication can be found at http://welikesharingdata.blob.core.windows.net/forshare/index.html.

### Authors’ contributions

EC designed and developed the software as well as wrote the paper. BAA and PA prepared all cell cultures and conducted the subsequent imaging. JH supervised both software and experimental design. All authors have read and approved the final manuscript.

## Acknowledgements

The authors would like to thank Paulos Lab for providing previously acquired mouse PMEL cell samples and Mehtora Lab for help with the cytospin. They would also like to thank Google for supporting our work through providing research credits (#38362142) for the Google Cloud Platform, which enables us to share our complete raw imaging data set with other scientists. Additionally, they thank River Abedon for his contributions to our early image segmentation and modeling efforts. This work is supported in part by the Flow Cytometry and Cell Sorting Unit Shared Resource, Hollings Cancer Center, Medical University of South Carolina (P30 CA138313).

## Footnotes

1 CODEX reference data was already aligned and deconvolved so segmentation was applied separately from preprocessing operations used for time benchmarks

